# Soaring migrants flexibly respond to sea-breeze in a migratory bottleneck: using first derivatives to identify behavioural adjustments over time

**DOI:** 10.1101/2022.12.15.520614

**Authors:** Paolo Becciu, David Troupin, Leonid Dinevich, Yossi Leshem, Nir Sapir

## Abstract

Millions of birds travel every year between Europe and Africa detouring ecological barriers and funnelling through migratory corridors where they face variable weather conditions. Little is known regarding the response of migrating birds to mesoscale meteorological processes during flight. Specifically, sea-breeze has a daily cycle that may directly influence the flight of diurnal migrants. We collected radar tracks of soaring migrants using modified weather radar in Latrun, central Israel, in 7 autumns between 2005 and 2016. We investigated how migrating soaring birds adjusted their flight speed and direction under the effects of daily sea-breeze circulation. We analysed the linear and, uniquely, the non-linear effects of wind on bird ground-, air- and sideways speed as function of time along the day using Generalized Additive Mixed Models and calculated first derivatives to identify when birds adjusted their response to the wind over time. Using data collected during a total of 148 days, we characterised the diel dynamics of horizontal wind flow in its two vectorial components relative to soaring migration goal (South), finding a consistent rotational movement of the wind blowing towards the East (morning) and to the South-East (late afternoon), with highest speed of crosswind component around mid-day and increasing tailwinds towards the late afternoon. We found that the airspeed of radar detected birds decreased consistently with increasing tailwind throughout the day, resulting in a rather stable groundspeed of 16-17 m/s. In addition, birds increased their sideways speed when crosswinds were at their maximum to an extent similar to that of the wind’s sideways component, meaning a full compensation to wind drift, which decreased after the time of crosswind maximum. Using a simple, novel and broadly applicable statistical method, we studied, for the first time, how wind influences bird flight by highlighting non-linear effects over time, providing new insights regarding the behavioural adjustments in the response of soaring birds to wind conditions. Our work enhances our understanding of how migrating birds respond to changing wind conditions during their journeys in order to exploit migratory corridors.

## Introduction

Geographic features and atmospheric conditions influence the movement of flying animals at different spatiotemporal scales [1, 2], yet little is known regarding the effects of mesoscale meteorological process on aerial flyers. Migrating birds, as well as other animals, may adjust their flight direction and speed in relation to weather and topography they encounter in order to accomplish their journey while saving time and energy, but these adjustments have only been seldom quantified. Importantly, the birds’ successful arrival at breeding or wintering grounds depends on their capacity to make space- and time-sensitive decisions to minimizing their energetic cost of travel and the total duration of migration [3, 4]. This can be achieved by avoiding hindering weather conditions (e.g., headwinds) and exploiting advantageous ones (e.g., tailwinds) encountered en route, which presumably induce fitness-related benefits [5, 6]. Soaring land migrants are often funnelling into so-called migratory bottlenecks and corridors, which are characterized by geographic features that allow them to minimize their cost of transport [7–9], compared to nearby areas [10]. When flying through a migratory corridor or bottleneck, birds are still at the mercy of weather. Notably, the migrants might be impacted by winds that may blow at various speeds and come from different directions in relation to the birds’ intended migration direction [11, 12]. Importantly, adverse winds, including head and lateral winds that may increase bird flight energetics, could slow the birds down or even terminate their flight [11, 13].

Soaring birds can avoid drifting off course by avoiding strong crosswinds and may reduce their cost of transport by preferentially migrating when favourable tailwinds prevail [14, 15]. However, if tailwinds are infrequent, waiting for ideal conditions could result in substantial delay of the journey [13, 16]. Initiating flight under crosswinds may result in drifting away from the intended track, with potentially severe consequences [17]. In-flight migrants may drift sideways due to crosswinds, or could try countering crosswinds by orienting towards the incoming wind, to compensate for the effect of crosswind fully or partially [11, 15, 18, 19]. Sideways drift compensation may diminish groundspeed, rendering it a suboptimal strategy [18]. Conversely, drifting birds may maximize groundspeed at the cost of geographic displacement, which may result in decreased fitness [17, 18]. Soaring migrants were found to flexibly change their flight speed and direction in relation to different wind conditions at different stages of their migration journey [15]. Specifically, Honey buzzards (*Pernis apivorus*) that crossed North-western Africa used different routes in autumn and spring; for instance, in autumn, they overcompensated for west-ward winds to circumvent the Atlas Mountains from their eastern side and then drifted with south-westward winds while crossing the Sahara Desert [15]. Probably, these different responses to wind allowed them to expend less energy and maximize their migration speed. Yet, many studies exploring the effects of wind on migrating birds involve between-day analyses that cannot usually portray within-day behavioural adjustments to dynamic wind conditions. In addition, studies of soaring birds usually involve either studying a rather small number of individuals over large spatial scales using tracking devices [18] or researching birds at specific locations [20]. Nevertheless, how huge waves of thousands of migrants adjust their flight to weather condition when flying through a certain migration corridor over several tens of kilometres is much less known. Furthermore, we still do not know if soaring migrants flexibly adjust their flight properties at different times of the day to maintain high migration speed and maintain their migration goal while facing dynamic wind patterns, such as sea breeze. The migration of soaring birds along the Mediterranean coast of the Levant region is massive and millions of soaring migrants are funnelled between the sea and the desert twice a year in a migration corridor parallel to the coastline [21, 22]. Along the coastline, especially in autumn, a dominant sea-breeze circulation process is characterized by a dynamic pattern through the course of the day with progressively increasing westerlies and clockwise change in wind direction towards the late afternoon and evening hours [23]. This process is created by the land-sea thermal gradient, and it is affected by several additional factors such as Coriolis force, large scale pressure gradients and friction (see [24] for a comprehensive review of this process). The sea-breeze front advances in a direction perpendicular to the coast towards inland and generates uplift with its vertical circulation component [23, 24]. The sea-breeze front has been hypothesised to influence the displacement of individual and flocks of birds in relation to the coastline. In Central Israel birds were observed to align to the sea-breeze front, supposedly exploiting the uplift to soar and increase their ground speed through the exploitation of this meteorologically dynamic process [25, 26]. The present study aims to examine how soaring birds are affected by the daily sea-breeze circulation during autumn migration, as well as their displacement relative to the coastline, using a modified weather radar in Central Israel. Specifically, we tested the following predictions: (a) if the wind rotates approximately from blowing eastwards to south-eastwards with the progression of the day, we expect that the birds will reduce their airspeed over time and concomitantly increase their groundspeed under the elevated tailwind conditions. Conversely, we predict that airspeed will increase, and groundspeed is expected to decrease under strong crosswind and headwinds [3, 4, 27–29]. (b) Wind drift compensation is predicted when winds are constant during the journey [30, 31], while changes in wind conditions are predicted to induce variable behaviour, such as compensation for crosswinds that may otherwise drift the birds towards the desert (east of the migration corridor) due to sea-breeze [11]. We aim to provide analytical framework for linking weather conditions to flight performance of diurnal migrants in an area that is characterized by intense soaring bird migration. This framework will allow integrating forecasted weather parameters to predict bird movement under a meso-scale meteorological dynamic process, namely sea-breeze circulation. We highlight the dynamic non-linear effects of the wind on bird flight and how bird response to the wind may change over time. Consequently, our work can help predicting bird movement and this information can be used for reducing the risk of collisions between birds and aircraft by separating aircrafts from birds in time and space [32].

## Methods

### Radar data

Data collection took place between 2005 and 2016 using a meteorological radar (MRL5) located in central Israel at Latrun (34.978^*◦*^N, 31.839^*◦*^E), 18 km southeast of the Ben Gurion International Airport (Fig. 1C). MRL5 is a two-wave meteorological radar, containing two simultaneously operating high-grade transmitters at two wavelengths (3 and 10 cm) and an antenna with two symmetrical narrow beams. The radius of the radar coverage area is 60 km (Fig. 1C). This radar is also equipped with two supplementary devices, a device for measuring echo signal fluctuations and a polarization device. This radar is the first meteorological radar adapted to exclusively detect birds with a series of algorithms that were applied to filter out other signals, such as those resulting from meteorological phenomena (i.e. clouds, rain, etc.) and human-related infrastructure and transportation (i.e. aircrafts, buildings, ships and ground vehicles). It furthermore classifies detected single birds into two main categories while providing their three-dimensional position and turning angle: local birds (low altitude and large turning angles) and migratory birds (higher altitudes and small turning angles). For a more detailed and technical explanation of the radar used in this study, please see [33–35]. We selected only radar data of diurnal autumn bird migration and specifically focused on the time window of Honey buzzard (*Pernis apivorus*) migration [22]. We consequently selected a period suitable for detecting Honey buzzard migration, between August 16^th^ and September 30^th^, but we note that during this period additional species migrated through the area (see below). We chose the Honey buzzard because it is the most abundant soaring migrant passing in the study area during this period, with more than 300,000 individuals counted on average in the autumn [22]. The other species of soaring migrants that pass through the area during this time of the year included black kites (*Milvus migrans*), Levant sparrowhawk (*Accipiter brevipes*), White stork (*Ciconia ciconia*) and Great white pelican (*Pelecanus onocrotalus*) [22, 36, 37]. Radar data included data collected from 3 hours after sunrise to 3 hours before sunset in order to consider only diurnal migration. We included data from the following 7 years (and days of data collection in each year): 2005 (29 days), 2006 (32 days), 2007 (32 days), 2009 (8 days), 2014 (17 days), 2015 (13 days) and 2016 (17 days) (Fig. S2). The skipped years in between contained fragmented and scarce data that were not used. The radar operated usually during several days in weekdays and never on weekends. Overall, data was analysed from a total of 148 days of radar operation. The radar was able to detect an echo of a bird and followed it in a 1.5-minute time window, collecting speed (m/s), altitude (m) and coordinates of the birds at the starting and ending points [35]. Every 15-30 minutes a .csv table was produced with the targets data along with a map image. The final dataset contained 857,206 tracks of diurnal soaring migrants.

**Figure 1:**
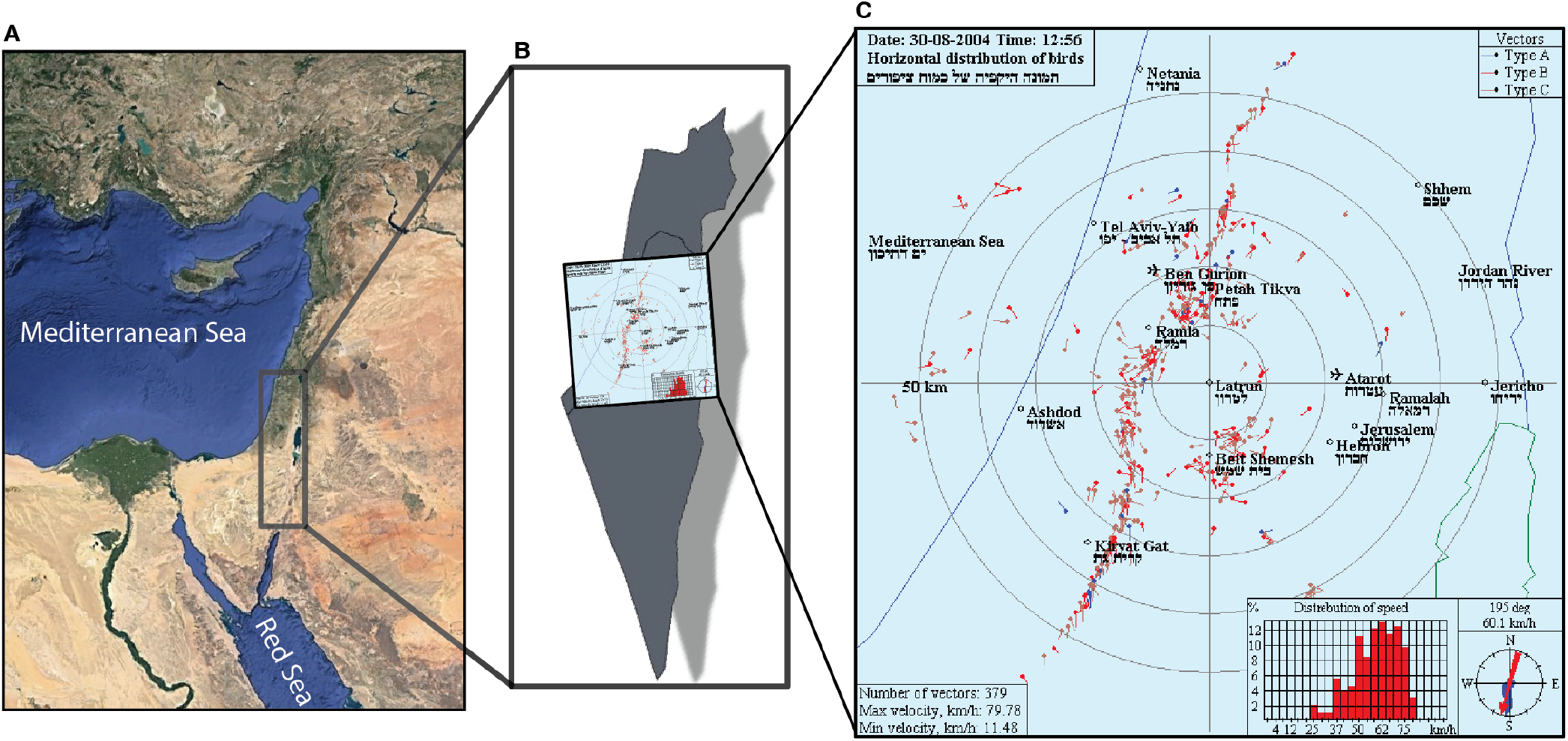
The study area located in the Levant region of the Middle East (A), where the radar is located in Central Israel (B), collecting bird tracking data between the Mediterranean Sea and the Jordan River (C).

### Weather data and movement parameters

We annotated the tracks with wind speed and direction from the closest weather station (Bet Dagan – 22 km west of the radar) of the Israeli Meteorological Service using a 10-minute interval database (available at: https://ims.data.gov.il). Wind data was collected at 10 m above the ground. The radar data was annotated with the closest wind data in time. We calculated tailwind and crosswind relative to the mean bird direction, simplified as 180 degrees (south) [38]. We calculated several flight parameters to explore the effects of the wind on bird movement during migration. We used the ground speed (*V*_*g*_ – distance covered in time in m/s) calculated for the trajectory between the initial and ending point recorded by the radar. We calculated the airspeed in m/s following Safi et al. (2013) [39]. To determine the birds’ lateral speed in m/s, we calculated a sideways speed with the following formula: *V*_*s*_ = *V*_*g*_ (*sin*(*θ* − π)), where *θ* is the track angle in radians [40].

### Data analysis

Bird tracks parameters, which include groundspeed, airspeed, sideways speed and the wind components, were averaged per recording session of the radar, every 15 minutes, except for 6.25% of the cases in which the time lag between consecutive sessions was 30 minutes. We clustered our 857,206 tracks into 1,977 sessions in which all the tracking parameters were averaged. These sessions constituted the final dataset for the analysis, to avoid temporal correlation of tracks recorded when conditions were similar for birds that flew at the same time within the radar coverage area. We used circular statistics (package circular [41] in the R environment [42]) such as the Rayleigh test to investigate the directionality of wind and bird directions. Then we fitted two GLMMs to model wind directions (in radians) and wind speed as a function of time to sunset (as quadratic term) in interaction with year as a grouping factor. We used a random slope structure with ordinal date as a random factor. We used the package function glmmTMB from the package glmmTMB [43] in R [42]. Furthermore, we fitted generalised additive mixed-effects models (GAMM) to assess how a) N-S and W-E components of the wind varied in their diurnal cycles (from 10 to 3 hours before sunset), if b) groundspeed and c) airspeed varied with the progression of the day following the daily wind pattern. Also, we used GAMM to check if the d) compensation-drift behaviour of the birds in relation to wind changed with the hour of the day. We used cubic regression penalized smoothing basis (k = 12) for the smooth term “hours before sunset” as a numeric continuous variable (for example a value of -6.25 means six hours and 15 minutes before the sunset), an autocorrelation-moving average correlation structure (corARMA, p = 2) for the “ordinal date”, and then used “year” and “ordinal date” as crossed random intercept. In order to run the GAMMs, we used the function gamm from the package mgcv [44] in R [42], with the “L-BFGS-B” non-linear optimization method for parameter estimation [45]. Furthermore, to identify the time of behavioural change, we calculated the first derivative *f ‘(x)*, highlighting significant periods of positive or negative relationships [46]. To do that, we used the function derivatives in gratia package [47]. The first derivative is the estimate *β* at each segment of the non-linear regression curve, which is the instantaneous rate of change of the function that defines the line. When the first derivative diverges from 0, there is a positive or negative response, evaluated as significant when the 95% simultaneous intervals (calculated at each step) are not overlapping with 0. We calculated the first derivative on a new set of predicted values (N = 1000, the number of points used for evaluating the derivative) from the GAMMs and calculated the simultaneous intervals using n_sim = 10,000 (the number of simulations used in computing the simultaneous intervals). As finite difference (eps) we set a value of 1e-07. We used simultaneous confidence intervals since they include information on model reliability. Hence, they are much more informative than pointwise confidence intervals to assess the goodness of a model that, subsequently, can be used to make inference or predictions [48, 49]. The code to run and plot the GAMM and its first derivative can be found in the online supplementary materials. Descriptive statistics reported in the text are mean *±* standard deviation, unless specified otherwise.

## Results

### The daily pattern of the wind during the migration period

Winds followed a consistent daily pattern during the study period showing a clockwise change of direction from the morning (about 10 hours before sunset) blowing towards 87.1 *±* 39.5 degrees, to the afternoon (about 3 hours before sunset) blowing towards 131.1 *±* 22.3 degrees. Mean wind direction was 112.5 *±* 26.7 degrees (Rayleigh test: *r* = 0.898, *P* < 0.0001, Fig.3B) with a mean wind speed of 4.1 *±* 1.1 m/s. The results of the GLMMs regarding wind direction (in radians) and speed as function of time among the years, highlighted that the wind direction changed slightly among years (see Table S1 and S2). More importantly, the wind direction rotates during the day in the same way among the years (Fig. 2A). Furthermore, wind speed did not change among the years (notably, 2005 had a slightly higher speed than the other years) and the increase in wind speed during the day followed a quadratic relationship with no differences among years (except for 2016, see Fig. 2B). The results of the GAMMs that examined the west-east and north-south components of the wind (crosswind and tailwind relative to the southward direction of soaring migrants, respectively) as function of hours before sunset, highlighted consistent daily wind patterns during the autumn among the 7 years included in the analysis. Crosswind speed increased from the beginning of the day (1 m/s), culminating in a plateau of 1-2 hours at the 7th and 6th hour before sunset (4 m/s), and then somewhat decreasing to 3 m/s towards the end of the day (Fig. 4A). This change in wind speed was stronger before the peak (with a rate of change ranging between 0.5 to about 1 m/s per hour), while after the peak, crosswind speed decreased at a stable rate of 0.3-0.4 m/s per hour (Fig. 4F). Tailwind had an almost linear relationship (*edf* = 1.86; Tab. S4) with the number of hours before sunset, with a consistently increasing speed throughout the day (Fig. 4B) at a stable rate of change of around 0.5 m/s per hour (Fig. 4G). These patterns demonstrate the shift in wind direction and speed throughout the day, with increasing wind speed and a shift between a current flowing primarily from west to east reaching a peak in wind speed at the middle of the day and rotating southward, resulting in a stronger tailwind component at the later hours of the day (see Fig.3).

**Figure 2:**
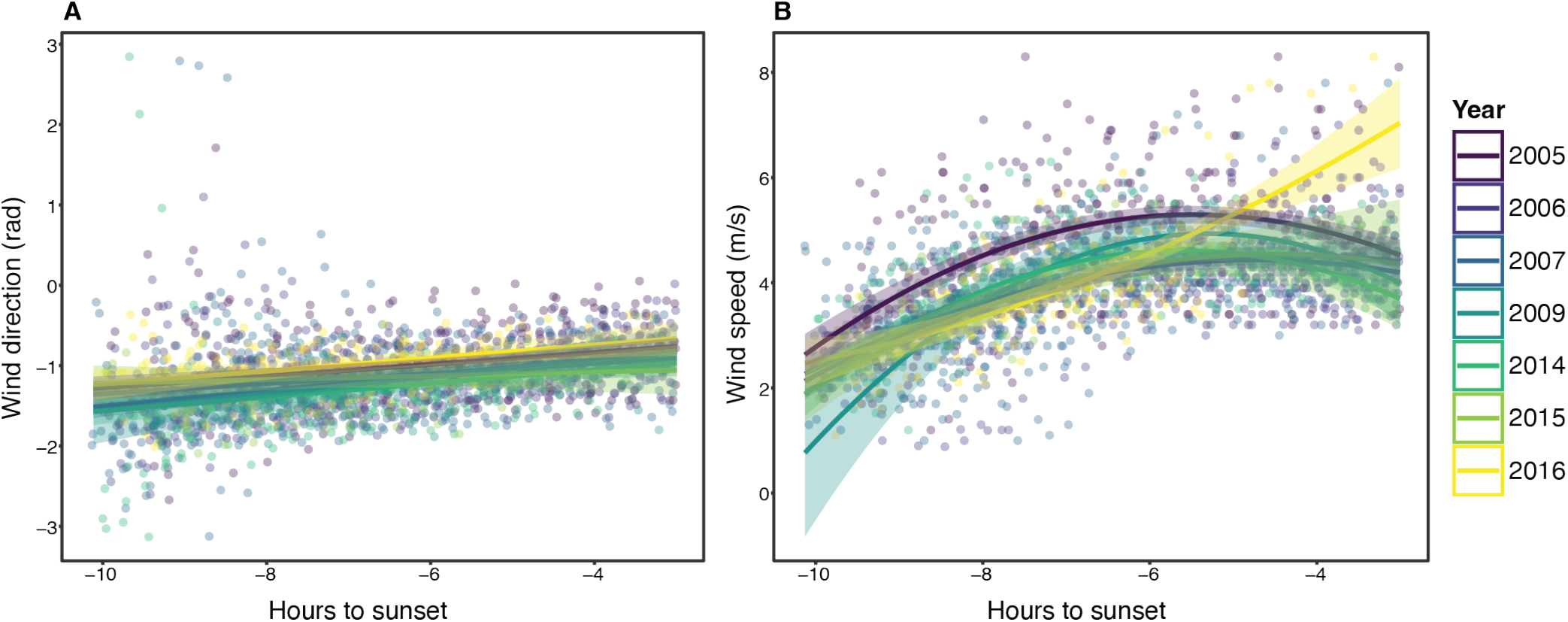
Predicted regression slopes (with 95% C.I.) of wind direction (A) and wind speed (B) predicted by the hour of the day relative to the sunset. Colours highlight the year of data collection.

**Figure 3:**
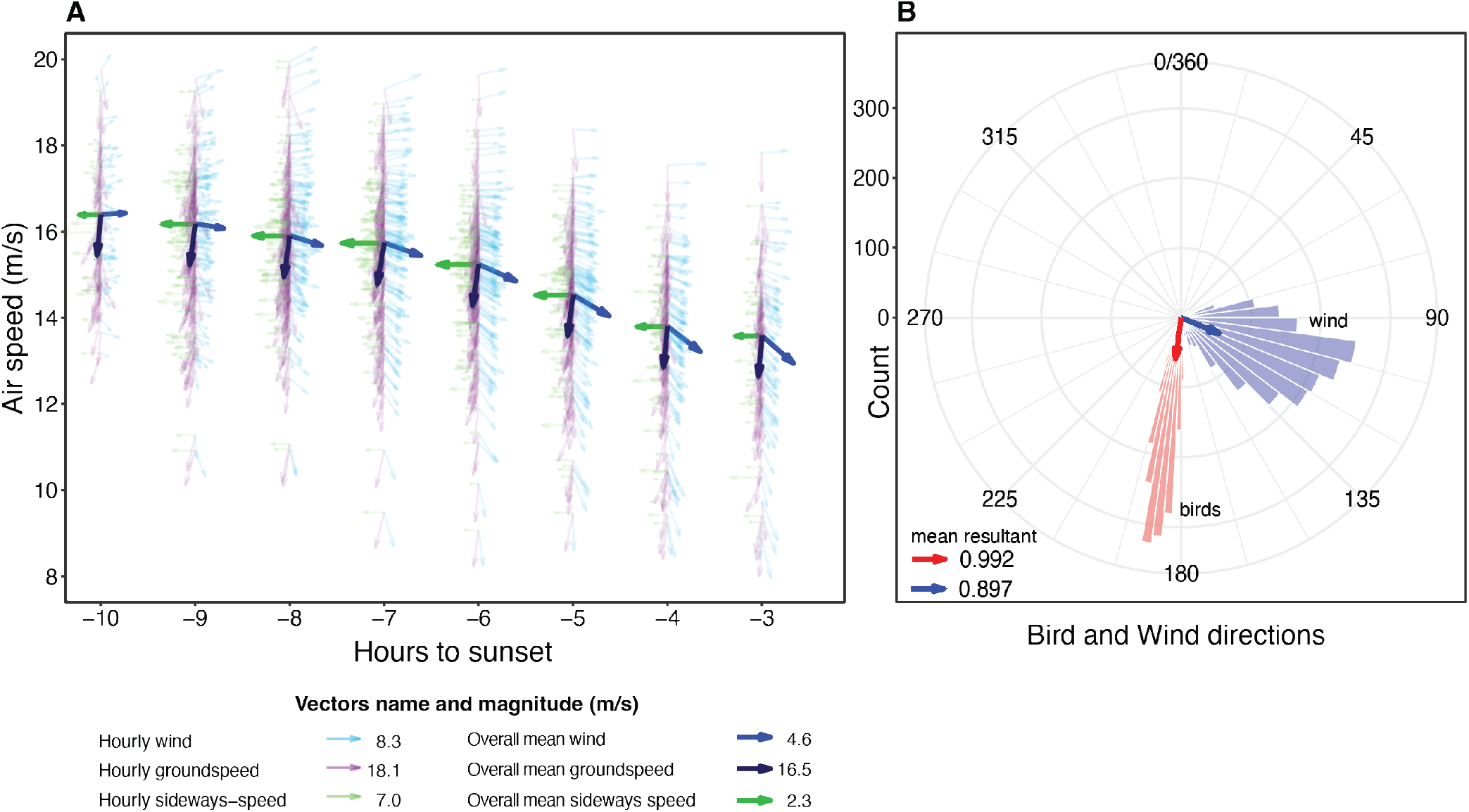
Visual representation of the flight directions and speeds of soaring migrants together with the wind conditions. Panel A shows that airspeed decreased throughout the day with overlapped vectors of mean groundspeed, sideways speed and wind speed and direction. Note that partially transparent values indicate the variation for each hour including data from each day of tracking from all the 7 years used in this study (2005-07, 2009, 2014-16). For visual purposes the vectors are not in scale, see the legend to contextualize the vector magnitude. Panel B shows circular histograms of wind directions (blue) and soaring bird flight directions (red), the arrows represent the mean vector and magnitude of the directions following Rayleigh tests (in which 0 = “uniform distribution” and 1 = “all the data are in the same direction”).

**Figure 4:**
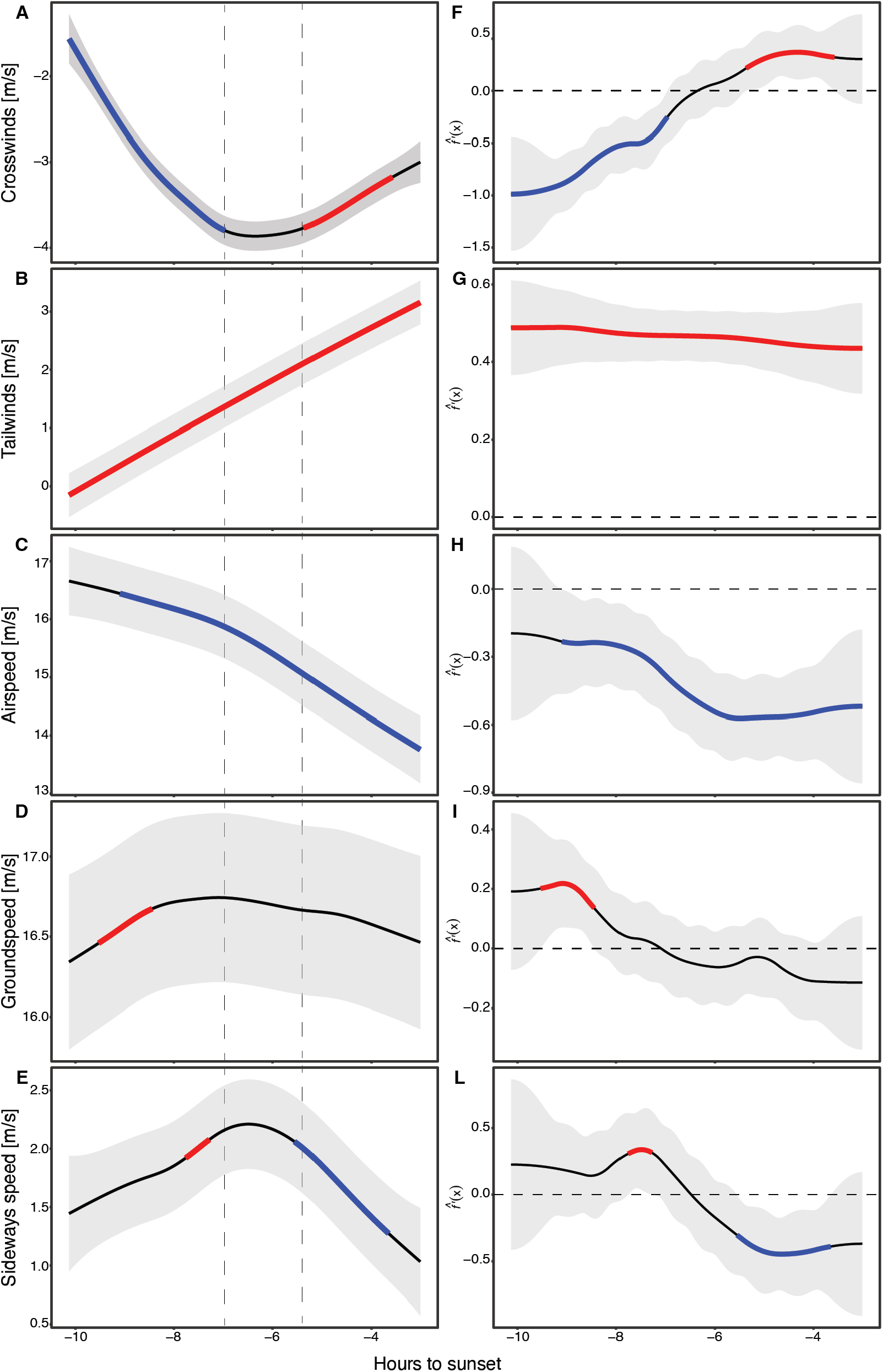
Cubic splines of the Generalized Additive Mixed Models with 95% C.I. (left side panels) and their first derivative estimations and 95% simultaneous confidence intervals (right side panels). Left panels: Estimated non-linear change of wind component and bird speeds (y-axis) as function of time (x-axis). Coloured sections indicate a significant rate of change that either increased (red) or decreased (blue) in that specific time period. The colouring of portions of the GAMM smoother line was adopted after calculating the first derivatives. Right panels: when the rate of change *f ‘(x)* in the y-axis is significantly above or below 0, the estimated rate of change is highlighted in blue (decrease) or red (increase).

### The daily pattern of movement of soaring migratory birds

Birds kept a highly consistent daily track direction of 187.7 *±* 0.1 degrees (Rayleigh test: *r* = 0.992, *P* < 0.0001) with a mean ground speed of 16.3 *±* 0.8 m/s (Fig. 3). Generalized Additive Mixed Models (GAMMs) and calculated first derivatives allowed us to compare dynamic behavioural change against wind change through time. This highlights, for the first time, periods and quantification of behavioural changes during diurnal migration in soaring migrants. We show that the birds’ airspeed decreased during the day (Fig. 4C) essentially at two main rates of change (Fig. 4H): in the first hours of the day when tailwinds were weak (< 1.5 m/s) the airspeed decreased at a rate of 0.2 m/s (Fig. 4H), while when the crosswind peak has passed and tailwind increased, airspeed decreased at a rate of 0.5 m/s (exactly opposite to the tailwind’s rate of change, Fig. 4G). Groundspeed was maintained between 16 and 17 m/s throughout the entire day (Fig. 4D), with a small change in the first two hours recorded (Fig. 4I). This change matched very low-speed winds and the birds’ highest airspeeds, probably also involving some flapping flight, which is usually not preferred by large soaring-gliding flyers. In addition, the birds’ sideways speed demonstrates a drift compensation behaviour relative to the encountered wind conditions (Fig. 3). The birds’ sideways speed remained above zero with a peak around the time of crosswind speed peak, and then decreased towards the end of the day (Fig. 4E). The pattern follows the opposite of that of crosswinds, meaning that birds dynamically compensated for crosswind throughout the entire day, to maintain a direction around 187 degrees (Fig. 3). This compensation changed during the day, increasing significantly at higher rate of change before the crosswind peak and decreasing significantly after it (Fig. 4L).

## Discussion

With an unprecedented amount of data of almost a million tracks from seven years, we quantitatively show how the flight of migrating soaring birds is predictable throughout the day and linked to a seasonal daily sea-breeze circulation pattern. We described the characteristics of the diel pattern of the wind flow encountered by the birds during the autumn season, coinciding with the migration period of Honey buzzards (*Pernis apivorous*) over a key corridor located between two ecological barriers for migrating birds: the eastern basin of the Mediterranean Sea and the Syrian Desert [21]. The diel circulation of the wind, consistently blowing to the east at the beginning of the day and shifting towards south-east at the end of the day, with an increasing speed, is apparently affecting bird response in relation to the wind. This response includes a reduction in soaring migrant airspeed with increasing tailwinds, which resulted in a rather constant groundspeed. Furthermore, the birds flew sideways, opposite the crosswinds, at a similar magnitude to their potential drift, suggesting full compensation when crosswind is the strongest. These flight responses to predictable weather conditions probably allow soaring birds to minimize their cost of transport (at least for most of the day) and keep a constant migration speed of around 16.5 m/s which would account for roughly 415 km (16.5 *·* 3.6 *·* 7, as [mean speed] x [km/h conversion constant] x [hours of migration considered here], respectively) covered during a single day in this section of their migration flyway [50]. By compensating for wind drift, the birds kept the migration direction towards the south-south-west, parallel to the coastline. We provide a methodological framework to study similar situations of non-linear relationships with the implementation of GAM(M)s through calculation of first derivatives with simulated simultaneous intervals (following the method suggested by Simpson [46]). This method mainly adds a fundamental step in the use of GAMMs in time series analysis, which is the calculation of first derivatives that highlight periods of change in the modelled non-linear relationships. This method is readapted here to study how bird response changes over time, representing the first application of this method in studies of animal behaviour after it was recently used in climate and paleoclimatic change studies [46, 51, 52]. This methodology allowed us to explore whether bird flight behaviour modification as the circulation of the wind varies over time. This method can be broadly applied to essentially any measurement that explores changes over time, including quantifying how different aspects of animal behaviour may change over time in relation to dynamic environmental conditions (i.e., temperature, wind, pressure, NDVI, etc.). Importantly, this method is not constrained by any time scale as long as the periodicity of the variable is accounted for by the GAMM. It may consequently be used for long (millennia) as well as short (minutes and seconds) time series. Furthermore, this method is easily applicable to highlight any behavioural change through time per se, or in relation to an event happening to the focal individual or population considered (e.g., predator-prey interaction, arrival to a specific location, inter- and intraspecific contacts, etc.). Along this line, the method proposed could be used to solve current challenges posed by (high-resolution) big data of animal movement [53, 54] or other fields of study. Our results confirm our first prediction regarding airspeed adjustment. As the wind rotated throughout the day, birds seemed to exploit the resultant increasing tailwind vector by decreasing their airspeed at a similar rate. Noteworthy, groundspeed did not increase with tailwinds but resulted in rather stable speed throughout the entire day. Hence, soaring migrants maintain a rather constant groundspeed under different winds. This may suggest that there is a cost for increased groundspeed – either a cost that relates to risks of grounding or switching to flapping flight that may bears metabolic expenses [20], or other possible costs associated with high groundspeed such as reduced flight control, hampered navigation and reduced sensing of subtle changes in airflow during flight, which might limit the use of convective updrafts for soaring. The soaring migrants’ sideways speed was always contrasting the crosswind that blew at different intensities towards the east (from the sea to the desert). To keep the migration direction and avoid wind drift, the birds compensated with dynamic modulation of their speed at opposing direction (see Fig. 4F and Fig. 4L). The increase in crosswind speed probably affected the birds’ airspeed. Consequently, the airspeed’s negative slope was substantially steeper after the peak of crosswinds speed, following by a concomitant decrease in sideways speed, when tailwind increased (see Fig. 4). This compensation for wind drift combined with a likely energy minimization strategy [4, 55, 56] allowed the soaring migrants to maintain the direction parallel to the coast at a constant and fairly high speed. Hence, this compensation strategy allowed avoiding drifting eastwards such that the birds will need to cross a wide sea later during their journey, after being drifted to South Sinai or even the Arabian Peninsula, as opposed to crossing the Suez Canal in North Sinai over a continuous landmass. Interestingly, by migrating parallel to the coastline, the birds fly over areas that contain trees, making the southern coastal plains of Israel and the northern part of Sinai the most suitable area for roosting before taking the leap over the Sahara Desert [22, 26]. This study was possible because of an exceptional adaptation of the MRL-5 weather radar, which was applied to detect only biological targets and filtered out weather and human-related features [33–35]. This allowed collecting an enormous quantity of bird migration data. Diurnal bird migration includes almost one million tracks and many more nocturnal migration tracks were recorded by this system. The collected data is unique since it isolates the target, locks on it for a certain time and allows creating a short track. This output is comparable (albeit producing shorter tracks) to those of modified marine radars or tracking radars, with the advantage of scanning 10 to 100 folds larger area. Therefore, the combination of having precise movement data of a huge number of targets for several years allowed us to undoubtedly quantify behavioural variation and quantify passage of flying migrants in one of the busiest migration corridors in the entire globe [57]. Importantly, this dataset may address additional questions regarding bird migration, providing extremely valuable information regarding the risk of bird strikes near several military and civil airports with heavy aircraft traffic. Our work provides important insights for understanding and predicting diurnal migration movements over an area where a significant number of collisions between aircrafts and birds take place every year (total of 402 bird strikes between 2000 and 2016), causing severe economic losses and risking human lives [32]. Knowing and predicting bird movement and response to wind conditions (i.e., ground-, sideways- and airspeed modulation relative to the predictable wind) may additionally be helpful for predicting movement not only over central Israel but also over other areas of the Levant region and elsewhere in the world, where migrating birds face similar sea-breeze circulations [58].

## Author contribution

P.B. and N.S. conceived the study, P.B. analysed the data, and wrote the first draft of this manuscript. Y.L and L.D. deployed and operated the radar, collected the radar data and maintained the database. All authors contributed to the final draft of the manuscript.

## Acknowledgements

We would like to thank the Israel Meteorological Service for providing the weather data.

