## Supplementary material for "Soaring migrants flexibly respond to sea-breeze in a migratory bottleneck: using first derivatives to identify behavioural adjustments over time": Fig. S1 Fig. S2 Fig. S3

### Electronic Supplementary Materials

Content:

- **Fig. S1 –** Radar tracks count and operating radar timeline
- **Fig. S3 –** Descriptive plots with data used for the GAMMs
- **Tables S1-S2 –** GLMM tables
- **Tables S3-S9 –** GAMM tables
- **Fig. S4 –** GAMM plots and first derivatives of 2 parameters not reported in the main text: “vertical speed” and “distance to the coast”

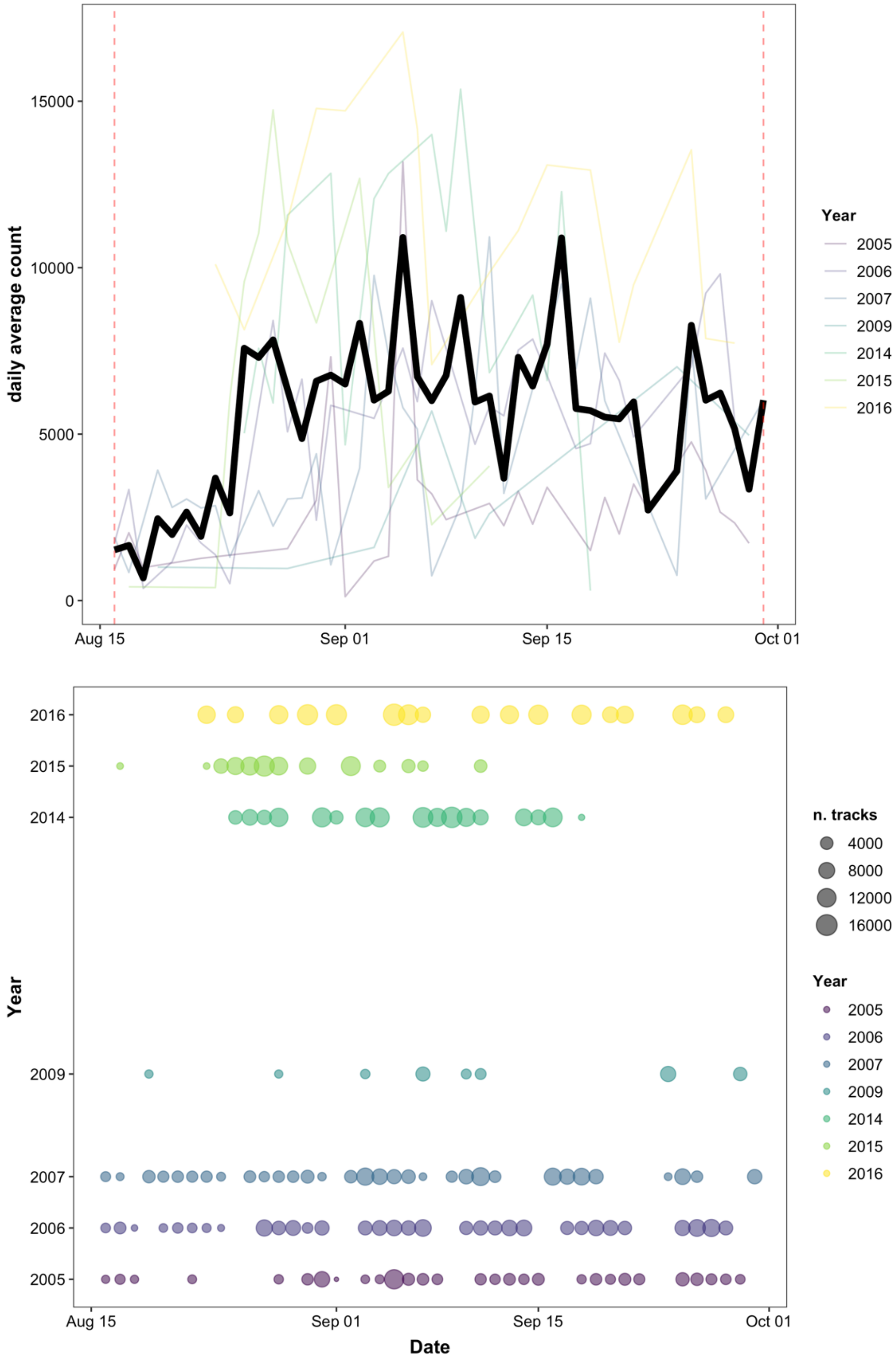

Fig. S1 – Top panel: Count of recorded radar tracks over the years considered for this study. The thicker black line is the average count of radar tracks per day in the years of the study (2005-07, 2009, 2014-16). The dashed lines delimit the time window considered for the Honey Buzzard migration (15 Aug - 30 Sep). Bottom panel: Operating days of the radar in the study period. Each circle corresponds to a day of the year, and the size is the sum of radar tracks recorded that day.

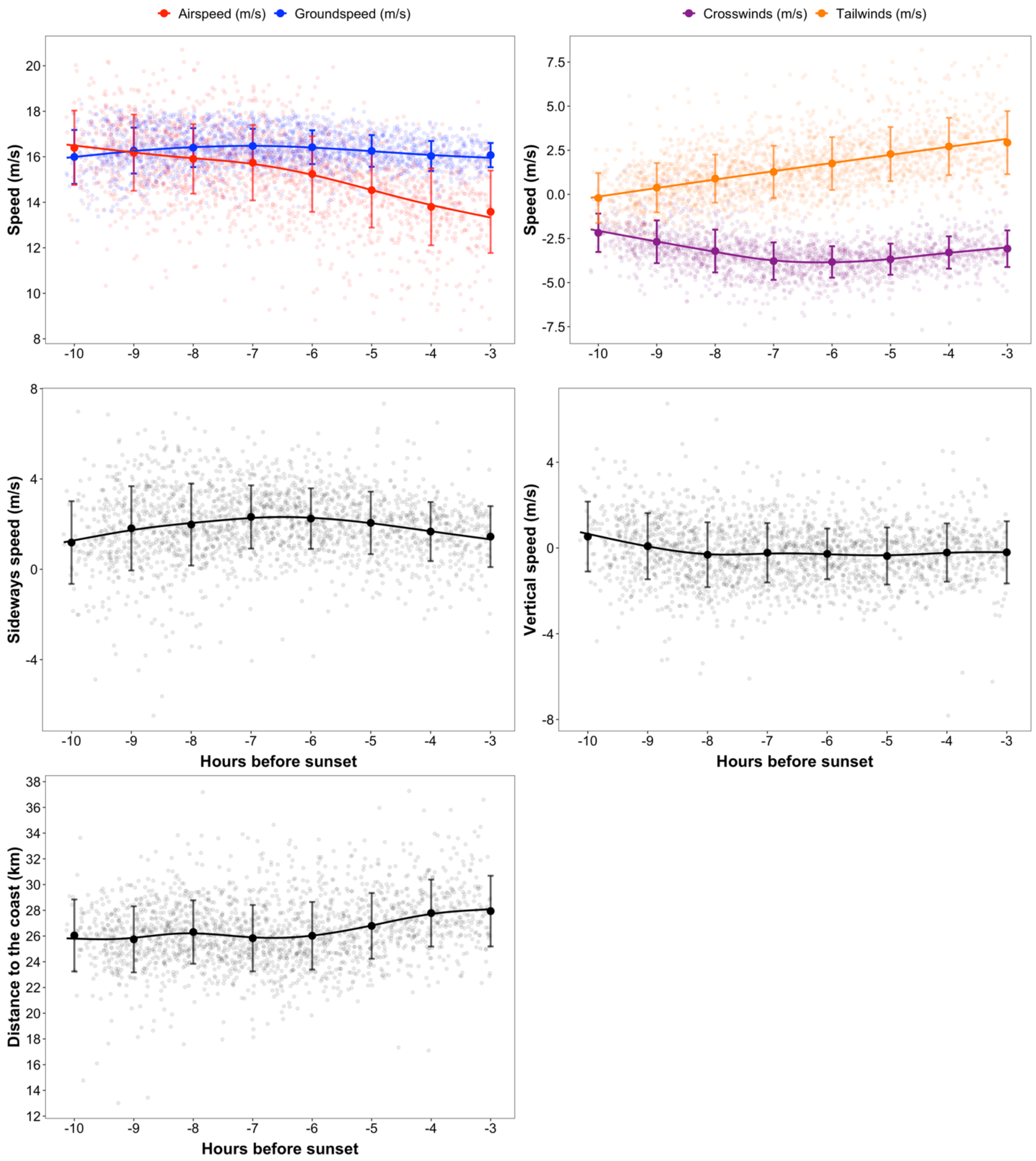

Fig. S3 – Set of graphs illustrating the data averaged per radar recording session (every 15 minutes, n = 1977 data points). bigger points are mean per hour with error bars showing standard deviation and a gam smoother highlighting the trend of the data.

#### Model tables

##### Wind direction GLMM table

| group | | Estimate | Standard Error | | statistic | p-value | |
|---|---|---|---|---|---|---|---|
| Fixed effects | | | | | | | |
| | (Intercept) | -1.055 | 0.038 | | -27.821 | 0.0000 | *** |
| | scale(h.from.sunset) | 0.145 | 0.019 | | 7.712 | 0.0000 | *** |
| | yearf2006 | -0.141 | 0.030 | | -4.643 | 0.0000 | *** |
| | yearf2007 | -0.102 | 0.033 | | -3.121 | 0.0018 | ** |
| | yearf2009 | -0.148 | 0.081 | | -1.823 | 0.0683 | . |
| | yearf2014 | -0.242 | 0.038 | | -6.320 | 0.0000 | *** |
| | yearf2015 | -0.064 | 0.057 | | -1.108 | 0.2677 | |
| | yearf2016 | 0.073 | 0.041 | | 1.759 | 0.0785 | . |
| Random effects | | | | | | | |
| n.days | sd__(Intercept) | 0.195 | | | | | |
| n.days | sd__scale(h.from.sunset) | 0.105 | | | | | |
| n.days | cor__(Intercept).scale(h.from.sunset) | -0.667 | | | | | |
| Residual | sd__Observation | 0.420 | | | | | |
| Signif. codes: 0 <= '***' < 0.001 < '**' < 0.01 < '*' < 0.05 | | | | | | | |

Tab. S1 – Summary table of the GLMM with wind direction in radians as response variable. The table reports the relationship between time of the day (hours before sunset), year and the changing wind direction (see Fig. 2A in main text). Ordinal date was included as random slope with hours before sunset.

##### Wind speed GLMM table

| group | | Estimate | Standard Error | | statistic | p-value | |
|---|---|---|---|---|---|---|---|
| Fixed effects | | | | | | | |
| | (Intercept) | 4.635 | 0.078 | | 59.686 | 0.0000 | *** |
| | scale(h.from.sunset) | -2.527 | 0.301 | | -8.386 | 0.0000 | *** |
| | yearf2006 | -0.804 | 0.054 | | -14.821 | 0.0000 | *** |
| | yearf2007 | -0.743 | 0.058 | | -12.725 | 0.0000 | *** |
| | yearf2009 | -0.736 | 0.167 | | -4.417 | 0.0000 | *** |

| group | | Estimate | Standard Error | | statistic | p-value | |
|---|---|---|---|---|---|---|---|
| | yearf2014 | -0.703 | 0.068 | | -10.282 | 0.0000 | *** |
| | yearf2015 | -0.805 | 0.127 | | -6.317 | 0.0000 | *** |
| | yearf2016 | -0.315 | 0.089 | | -3.545 | 0.0004 | *** |
| | I(scale(h.from.sunset^2)) | -3.039 | 0.306 | | -9.934 | 0.0000 | *** |
| | scale(h.from.sunset):yearf2006 | 1.135 | 0.371 | | 3.055 | 0.0022 | ** |
| | scale(h.from.sunset):yearf2007 | 1.464 | 0.383 | | 3.823 | 0.0001 | *** |
| | scale(h.from.sunset):yearf2009 | -0.987 | 1.119 | | -0.882 | 0.3776 | |
| | scale(h.from.sunset):yearf2014 | -0.263 | 0.473 | | -0.556 | 0.5780 | |
| | scale(h.from.sunset):yearf2015 | 1.117 | 0.971 | | 1.151 | 0.2499 | |
| | scale(h.from.sunset):yearf2016 | 4.813 | 0.743 | | 6.475 | 0.0000 | *** |
| | yearf2006:I(scale(h.from.sunset^2)) | 1.097 | 0.381 | | 2.880 | 0.0040 | ** |
| | yearf2007:I(scale(h.from.sunset^2)) | 1.410 | 0.386 | | 3.655 | 0.0003 | *** |
| | yearf2009:I(scale(h.from.sunset^2)) | -1.342 | 1.236 | | -1.086 | 0.2777 | |
| | yearf2014:I(scale(h.from.sunset^2)) | -0.247 | 0.472 | | -0.525 | 0.5997 | |
| | yearf2015:I(scale(h.from.sunset^2)) | 0.976 | 0.874 | | 1.117 | 0.2639 | |
| | yearf2016:I(scale(h.from.sunset^2)) | 4.147 | 0.693 | | 5.982 | 0.0000 | *** |
| Random effects | | | | | | | |
| n.days | sd__(Intercept) | 0.431 | | | | | |
| n.days | sd__scale(h.from.sunset) | 0.293 | | | | | |
| n.days | cor__(Intercept).scale(h.from.sunset) | -0.423 | | | | | |
| Residual | sd__Observation | 0.736 | | | | | |
| Signif. codes: 0 <= '***' < 0.001 < '**' < 0.01 < '*' < 0.05 | | | | | | | |

Tab. S2 – Summary table of the GLMM with wind speed (m/s) as response variable. The table reports the relationship between time of the day (hours before sunset) and its quadratic term in interaction with the year and the changing wind speed (see Fig. 2B in main text). Ordinal date was included as random slope with hours before sunset.

#### Crosswinds GAMM table

| Component | Term | Estimate | Std Error | t-value | p-value |
|---|---|---|---|---|---|

| Component | Term | Estimate | Std Error | t-value | p-value | |
|---|---|---|---|---|---|---|
| A. parametric coefficients | (Intercept) | -3.344 | 0.068 | -48.911 | < 0.0001 | *** |
| Component | Term | edf | Ref. df | F-value | p-value | |
| B. smooth terms | s(h.from.sunset) | 6.617 | 6.617 | 48.464 | < 0.0001 | *** |
| Signif. codes: 0 <= '***' < 0.001 < '**' < 0.01 < '*' < 0.05 | | | | | | |
| Adjusted R-squared: 0.158 | | | | | | |
| N: 1977 | | | | | | |

Tab. S3 – Summary table of the GAMM with crosswind component of the wind as response variable. The table reports the intercept and the complexity of the fitted cubic spline (edf).

#### Tailwinds GAMM table

| Component | Term | Estimate | Std Error | t-value | p-value | |
|---|---|---|---|---|---|---|
| A. parametric coefficients | (Intercept) | 1.481 | 0.186 | 7.943 | < 0.0001 | *** |
| Component | Term | edf | Ref. df | F-value | p-value | |
| B. smooth terms | s(h.from.sunset) | 1.858 | 1.858 | 184.181 | < 0.0001 | *** |
| Signif. codes: 0 <= '***' < 0.001 < '**' < 0.01 < '*' < 0.05 | | | | | | |
| Adjusted R-squared: 0.253 | | | | | | |
| N: 1977 | | | | | | |

Tab. S4 – Summary table of the GAMM with tailwind component of the wind as response variable. The table reports the intercept and the complexity of the fitted cubic spline (edf).

#### Airspeed GAMM table

| Component | Term | Estimate | Std Error | t-value | p-value | |
|---|---|---|---|---|---|---|
| A. parametric coefficients | (Intercept) | 15.563 | 0.285 | 54.667 | < 0.0001 | *** |
| Component | Term | edf | Ref. df | F-value | p-value | |
| B. smooth terms | s(h.from.sunset) | 3.728 | 3.728 | 62.854 | < 0.0001 | *** |
| Signif. codes: 0 <= '***' < 0.001 < '**' < 0.01 < '*' < 0.05 | | | | | | |
| Adjusted R-squared: 0.216 | | | | | | |
| N: 1977 | | | | | | |

Tab. S5 – Summary table of the GAMM with bird airspeed as response variable. The table reports the intercept and the complexity of the fitted cubic spline (edf).

#### Groundspeed GAMM table

| Component | Term | Estimate | Std Error | t-value | p-value | |
|---|---|---|---|---|---|---|
| A. parametric coefficients | (Intercept) | 16.648 | 0.267 | 62.245 | < 0.0001 | *** |
| **Component** | **Term** | **edf** | **Ref. df** | **F-value** | **p-value** | |
| B. smooth terms | s(h.from.sunset) | 4.940 | 4.940 | 6.771 | < 0.0001 | *** |
| Signif. codes: 0 <= '***' < 0.001 < '**' < 0.01 < '*' < 0.05 | | | | | | |
| Adjusted R-squared: 0.025 | | | | | | |
| N: 1977 | | | | | | |

Tab. S6 – Summary table of the GAMM with bird groundspeed as response variable. The table reports the intercept and the complexity of the fitted cubic spline (edf).

#### Sideways speed GAMM table

| Component | Term | Estimate | Std Error | t-value | p-value | |
|---|---|---|---|---|---|---|
| A. parametric coefficients | (Intercept) | 1.822 | 0.184 | 9.889 | < 0.0001 | *** |
| **Component** | **Term** | **edf** | **Ref. df** | **F-value** | **p-value** | |
| B. smooth terms | s(h.from.sunset) | 5.224 | 5.224 | 8.999 | < 0.0001 | *** |
| Signif. codes: 0 <= '***' < 0.001 < '**' < 0.01 < '*' < 0.05 | | | | | | |
| Adjusted R-squared: 0.0314 | | | | | | |
| N: 1977 | | | | | | |

Tab. S7 – Summary table of the GAMM with bird  sideways speed as response variable. The table reports the intercept and the complexity of the fitted cubic spline (edf).

#### Vertical speed GAMM table

| Component | Term | Estimate | Std Error | t-value | p-value | |
|---|---|---|---|---|---|---|
| A. parametric coefficients | (Intercept) | -0.139 | 0.164 | -0.849 | 0.3959 | |
| **Component** | **Term** | **edf** | **Ref. df** | **F-value** | **p-value** | |
| B. smooth terms | s(h.from.sunset) | 4.415 | 4.415 | 4.638 | 0.0005 | *** |
| Signif. codes: 0 <= '***' < 0.001 < '**' < 0.01 < '*' < 0.05 | | | | | | |
| Adjusted R-squared: 0.0188 | | | | | | |
| N: 1977 | | | | | | |

Tab. S8 – Summary table of the GAMM with bird vertical speed as response variable. The table reports the intercept and the complexity of the fitted cubic spline (edf).

#### Distance to the coast GAMM table

| Component | Term | Estimate | Std Error | t-value | p-value | |
|---|---|---|---|---|---|---|
| A. parametric coefficients | (Intercept) | 26.333 | 0.276 | 95.366 | 0.0000 | *** |
| **Component** | **Term** | **edf** | **Ref. df** | **F-value** | **p-value** | |
| B. smooth terms | s(h.from.sunset) | 6.771 | 6.771 | 14.271 | 0.0000 | *** |
| Signif. codes: 0 <= '***' < 0.001 < '**' < 0.01 < '*' < 0.05 | | | | | | |
| Adjusted R-squared: 0.064 | | | | | | |
| N: 1977 | | | | | | |

Tab. S9 – Summary table of the GAMM with bird distance to the coast as response variable. The table reports the intercept and the complexity of the fitted cubic spline (edf).

#### Panel figure – Vertical speed and Distance to the coast GAMMS and first derivatives

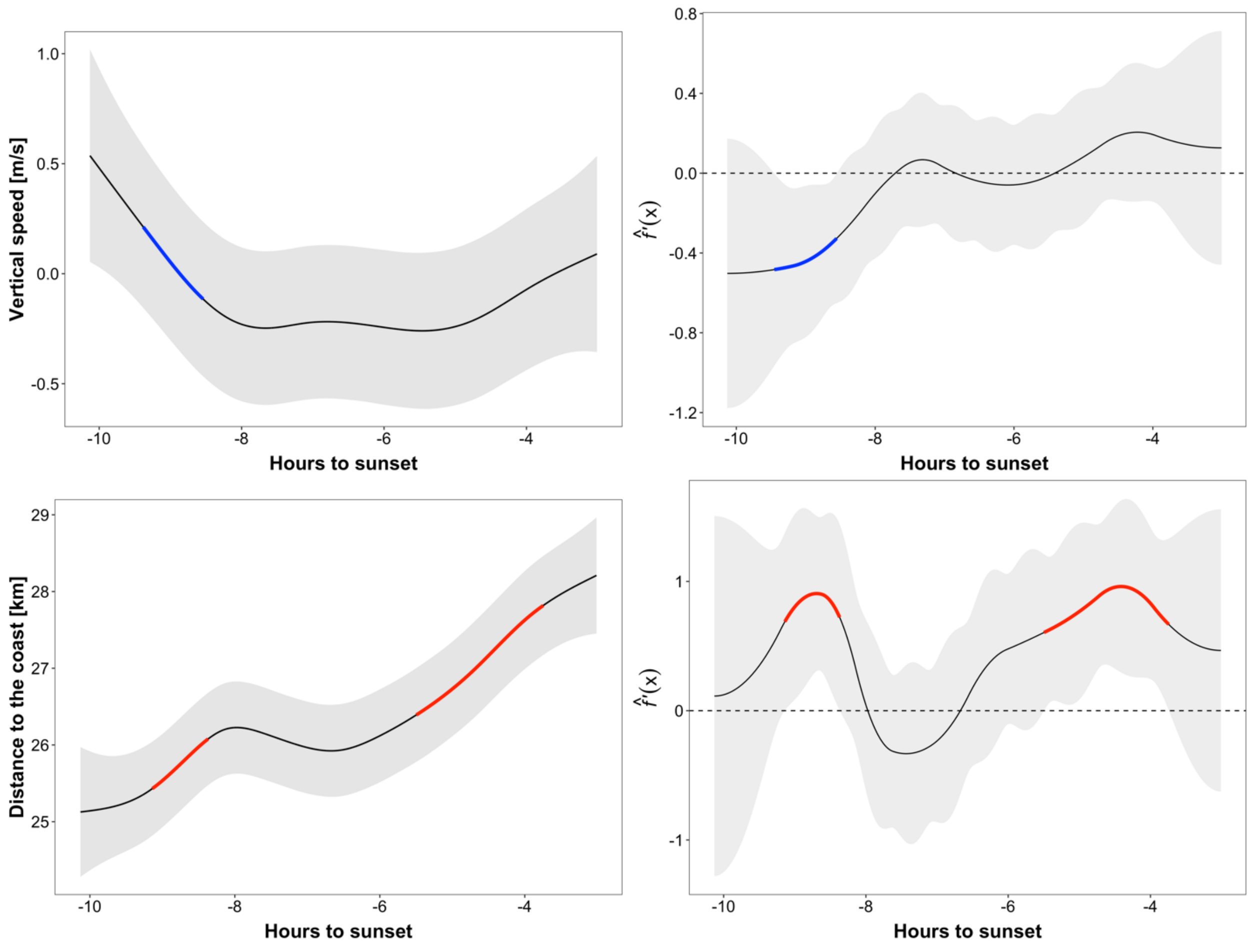

Fig. S4 – Cubic splines of the Generalized Additive Mixed Models with 95% C.I. (left side panels) and their first derivative estimations and 95% simultaneous confidence intervals (right side panels). Left panels: Estimated non-linear change of vertical speed and distance to the coast (y-axis) as function of time (x-axis), coloured portions depend on the rate of change that significantly increased (red) or decreased (blue) in that specific time period. The colouring of portions of the GAMM smoother line was adopted after calculating the first derivatives. Right panels: when the rate of change f’(x) in the y-axis is above or below 0 (including the 95% C.I.) the estimated rate of change is highlighted in blue (decrease) or red (increase).
